# AI-Powered Smart Glasses for Sensing and Recognition of Human-Robot Walking Environments

**DOI:** 10.1101/2023.10.24.563804

**Authors:** Daniel Rossos, Alex Mihailidis, Brokoslaw Laschowski

**Affiliations:** Division of Engineering Science and the Robotics Institute, University of Toronto, Toronto, Canada; KITE Research Institute, Toronto Rehabilitation Institute, Toronto, Canada; Institute of Biomedical Engineering and the Robotics Institute, University of Toronto, Toronto, Canada; Department of Mechanical and Industrial Engineering and the Robotics Institute, University of Toronto, Toronto, Canada

## Abstract

Environment sensing and recognition can allow humans and robots to dynamically adapt to different walking terrains. However, fast and accurate visual perception is challenging, especially on embedded devices with limited computational resources. The purpose of this study was to develop a novel pair of AI-powered smart glasses for onboard sensing and recognition of human-robot walking environments with high accuracy and low latency. We used a Raspberry Pi Pico microcontroller and an ArduCam HM0360 low-power camera, both of which interface with the eyeglass frames using 3D-printed mounts that we custom-designed. We trained and optimized a lightweight and efficient convolutional neural network using a MobileNetV1 backbone to classify the walking terrain as either indoor surfaces, outdoor surfaces (grass and dirt), or outdoor surfaces (paved) using over 62,500 egocentric images that we adapted and manually labelled from the Meta Ego4D dataset. We then compiled and deployed our deep learning model using TensorFlow Lite Micro and post-training quantization to create a minimized byte array model of size 0.31MB. Our system was able to accurately predict complex walking environments with 93.6% classification accuracy and had an embedded inference speed of 1.5 seconds during online experiments using the integrated camera and microcontroller. Our AI-powered smart glasses open new opportunities for visual perception of human-robot walking environments where embedded inference and a low form factor is required. Future research will focus on improving the onboard inference speed and miniaturization of the mechatronic components.

## 1 Introduction

Visual sensing and recognition of human-robot walking environments is of growing interest. Applications range from autonomous control and planning of robotic leg prostheses and exoskeletons to providing sensory feedback to persons with visual impairments. However, most egocentric visual perception systems, such as Project Aria by Meta [1], have been limited to off-device inferencing with external machines and cloud computing.

Previous research has mainly focused on head-mounted cameras with large computation systems such as Raspberry Pi 3 [2], [3], and chest and waist-mounted cameras [4]-[5]. However, these systems did not integrate the camera sensor and computation all within a single system. Other studies [6]-[11] have used large convolutional neural networks (CNNs) with many learnable parameters for accurate image classification of walking terrains, including level-ground, stairs, and other obstacles, though lacked onboard inferencing. Some researchers have used classical machine learning methods with success [12]-[16]. While both deep learning and non-deep learning models have shown good accuracy performance, they have not focused on deployment and efficiency as the main objectives. Consequently, these systems have been restricted to highpower computers and have limited deployment on mobile and embedded devices.

An integrated visual perception system has yet to be designed, prototyped, and evaluated on edge devices with low inference speeds. This gap could be explained by limitations in mobile and embedded computing, which have only recently been alleviated by advances in hardware and deep learning model compression methods. Accordingly, the purpose of this study was to develop AI-powered smart glasses that uniquely integrate both sensing and deep learning computation for visual perception of human-robot walking environments while achieving high accuracy and low latency (Fig. 1). We integrated our mechatronic components all within a single device, which is lightweight and a small form factor as to not obstruct mobility or user comfort. Computationally, it has sufficient memory and processing power for real-time inferencing with live video stream.

**Fig. 1.**
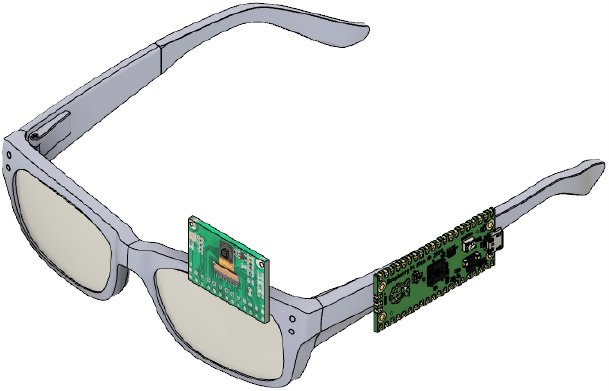
Computer design of our smart glasses, including the 3D-printed mechanical mounts used to interface the mechatronic components with the eyeglass frames.

## II. Methods

### A. Mechatronics Design

The mechanical mounts for our smart glasses included two design considerations: 1) the location of the mechatronic components on the frames, and 2) the means by which these components are attached. The location of the microcontroller and camera was partially inspired by commercial smart glasses such as Google Glass [17] and Ray-Ban Stories [18], which have the camera forward facing and the computational processor on the arms of the frames [19]. This design allows for a larger processor to not obstruct the visual field-of-view while also having the camera simulate the orientation and perspective of the user i.e., egocentric. We designed a semi-permanent mounting system that would allow our smart glasses to be applicable and transferable to a wide range of eyeglass frames. We custom-designed and 3D-printed mounting brackets for the camera and microcontroller (Fig. 2). The two main mechatronic components required to develop our system is the camera to capture visual information about the walking environment, and the microcontroller for processing and computing the images (Fig. 3). With the low-power, low-latency constraints of our design, heightened scrutiny of relevant metrics and constraints was required to identify optimal components.

**Fig. 2.**
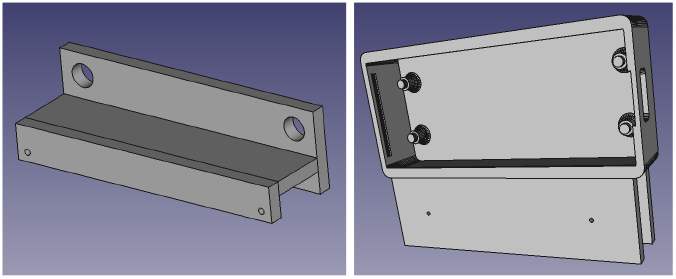
Our custom-designed 3D-printed mounts for the camera (left) and microcontroller (right) with screw holes to secure the mechatronics to the eyeglass frames using rubber-tipped screws.

**Fig. 3.**
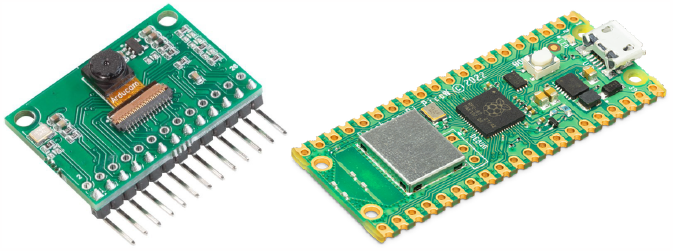
The Arducam HM0360 low-power camera (left) used for vision sensing and the Raspberry Pi Pico microcontroller (right) used for the onboard image processing and computation.

We used the ArduCam HM0360 VGA SPI camera due to its low power consumption, high frame rate, and high resolution [20]. The camera has a power consumption of less than 19.6mW during active VGA sampling. This low-power consumption supports the “always-on” operating mode that our smart glasses aim for by ensuring that the power consumption would be minimally affected by continuous sensing. Another important feature of the camera is the high frame rate. At 60 frames per second, this provides the microcontroller with a high enough sampling rate to ensure that there are no bottlenecks to the image classification resulting from the camera’s framerate, while also providing updated real-world visual data, reducing lag in our smart glasses’ understanding. The camera includes a high resolution of 640x480 monochrome images. This resolution provides an image size large enough to portray environmental state information and even supports down-sampling of the resolution to allow for smaller models and faster inference predictions.

For the onboard computational processing, we used the Raspberry Pi Pico W microcontroller. This newly developed board has more memory and CPU power compared to smaller boards. The increase in processing power and memory, small form factor, and capability for wireless communication made this embedded microcontroller a viable solution for our smart glasses. Compared to microcontrollers of comparable size, such as the Arduino Nano 33 BLE with a processing speed of 64 MHz, the Pico contains Dual ARM 133 MHz processors. This added processing power provides sufficient speed and parallelization to process live video streams while minimizing inference speeds, aligned with our goal of achieving real-time predictions using deep learning.

The Pico contains 64 kB SRAM and 2 MB QSPI flash memory, which is greater than other microcontrollers such as the Arduino Nano 33 BLE. This increased memory is important for running machine learning algorithms directly on the embedded device and providing flexibility in the type of models that can be loaded, including more memory-intensive models such as deep convolutional neural networks. The Pico also has a small form factor of 21mm x 51.3 mm, which is essential for our design to be easily integrated into eyeglass frames and provide minimal obstruction to mobility or user comfort. The Pico can wirelessly communicate and interface with external robotic devices and computers via a CYW43439 chip, which supports single-band 2.4 GHz Wi-Fi connection and Bluetooth 5.2.

### B. Computer Vision and Deep Learning

We created a new image dataset based on the Meta Ego4D dataset [21]. The full Ego4D dataset includes more than 3,670 hours of egocentric (i.e., first-person) video collected by 923 subjects from 74 locations worldwide (Fig. 4). The images were collected using head-mounted wearable cameras, which made the dataset highly applicable to our computer vision application. In addition to an appropriate camera angle, the video clips were pre-labelled to identify the scene in which the videos were recorded.

**Fig. 4.**
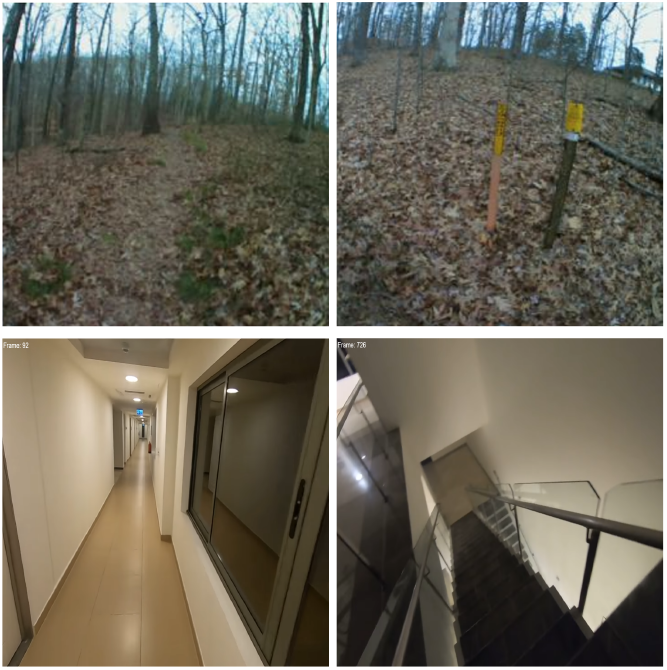
Examples of images we adapted and manually labelled from the Meta Ego4D dataset [21].

Extending the Ego4D annotations, we developed new class definitions and manually re-labelled images as either 1) indoor surfaces, 2) outdoor surfaces (grass and dirt), and 3) outdoor surfaces (paved). See Table 1 for our class distributions. We sampled the videos at one frame per second to collect images. To reduce the required memory storage to run our deep learning model, images in our dataset were downsampled to 96x96 pixels before being used for training, therein minimizing the staging area requirements for our microcontroller. This down sampling is a constraint imposed by the computing hardware. To help reduce overfitting during the model training, we added random horizontal reflections, image zooms, slight rotations, and contrast changes. These augmentations were deemed appropriate as such effects would likely occur in real-world walking while wearing glasses. Finally, all images were converted to grayscale to mimic the conditions of our onboard camera (Fig. 5).

**Table 1.** Breakdown of the class distributions in our new image dataset that we developed from the Meta Ego4D dataset [21].

| Environment class | Total frames | Number of videos | Frames per video |
| --- | --- | --- | --- |
| Indoor surfaces | 24,189 | 51 | 474 |
| Outdoor surfaces (paved) | 21,362 | 51 | 418 |
| Outdoor surfaces (grass and dirt) | 16,958 | 31 | 547 |

**Fig. 5.**
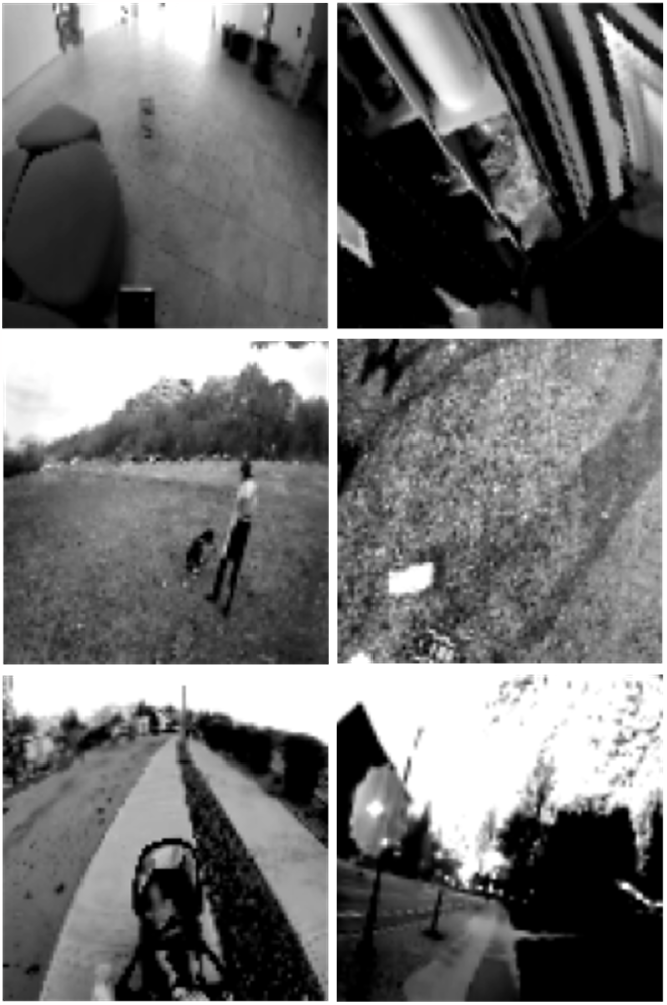
Examples of images from our glasses, including indoor surfaces (top row), outdoor surfaces - grass and dirt (middle row), and outdoor surfaces - paved (bottom row).

For image classification and automatic feature engineering, we used the base model of MobileNetV1 [22] in TensorFlow with an alpha value of 0.25, thereby reducing the model width and learnable parameters to lower the computational demand on our embedded device. An additional 2D convolutional layer was added before the MobileNet base model to expand the input dimensions of the grayscale images to a 3-channel image as required by the MobileNet layer. The MobileNet layer is followed by a 2D global average pooling layer to reduce the dimensionality of the 2D output, followed by a fully connected layer with a softmax activation to predict the three walking terrains (Table 2). The MobileNet architecture was selected as the underlying model similar to [7] because the depth-wise separable convolutional layers aid in efficient and accurate image classification. The model contained ∼219,300 parameters and was trained using TensorFlow. The dataset was split into training (70%), validation (15%), and test (15%) sets. To avoid data leakage between test and validation sets, the source videos for frames within the training and validation sets were different.

**Table 2.** The lightweight and efficient deep learning model [22] used for image classification of walking environment.

| Network layer | Output shape | Number of parameters |
| --- | --- | --- |
| Convolutional 2D | 1x96x96x3 | 30 |
| MobileNet_V1_0.25 | 1x3x3x256 | 218,544 |
| Global average pooling | 1x256 | 0 |
| Fully connected | 1x3 | 771 |

Finally, we converted our deep learning model to a TensorFlow Lite model using a quantization method converting the floating-point numbers to 8-bit integers and resolving incompatible tensor operations. The TensorFlow Lite model was then converted using the TensorFlow Lite Micro tooling to produce a byte array usable by the microcontroller for onboard inference. Our final model was of size 0.31MB. To quantify inference speed, we took the most recent image from the camera and loaded it to the microcontroller’s memory. The image was then loaded into memory as input to the model, and the resulting label for that frame was derived.

## III. Results

Our deep learning model achieved a training accuracy of 97.7%, a training loss of 0.07, a validation accuracy of 93.2%, and a validation loss of 0.41 (Fig. 6). During inference on the test set, the compressed model achieved an overall prediction accuracy of 93.6%, an f1-score of 93.6%, a precision of 93.7%, and a recall of 93.6%. The multiclass confusion matrix in Table 3 shows the distribution of the prediction accuracies for each walking environment. The neural network most accurately predicted outdoor surfaces - grass and dirt (96.8%), followed by outdoor surfaces - paved (94.7%) and indoor surfaces (90%). The onboard inference speed on the embedded device was 1.5 seconds from reading the image to outputting the predicted label.

**Table 3.**
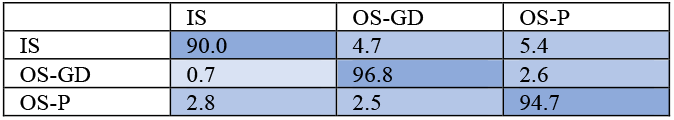
Multiclass confusion matrix showing the image classification accuracies (%) during inference on the test set. The columns and rows are the predicted and labelled classes, respectively. The classes include indoor surfaces (IS), outdoor surfaces - grass and dirt (OS-GD), and out- door surfaces - paved (OS-P).

**Fig. 6.**
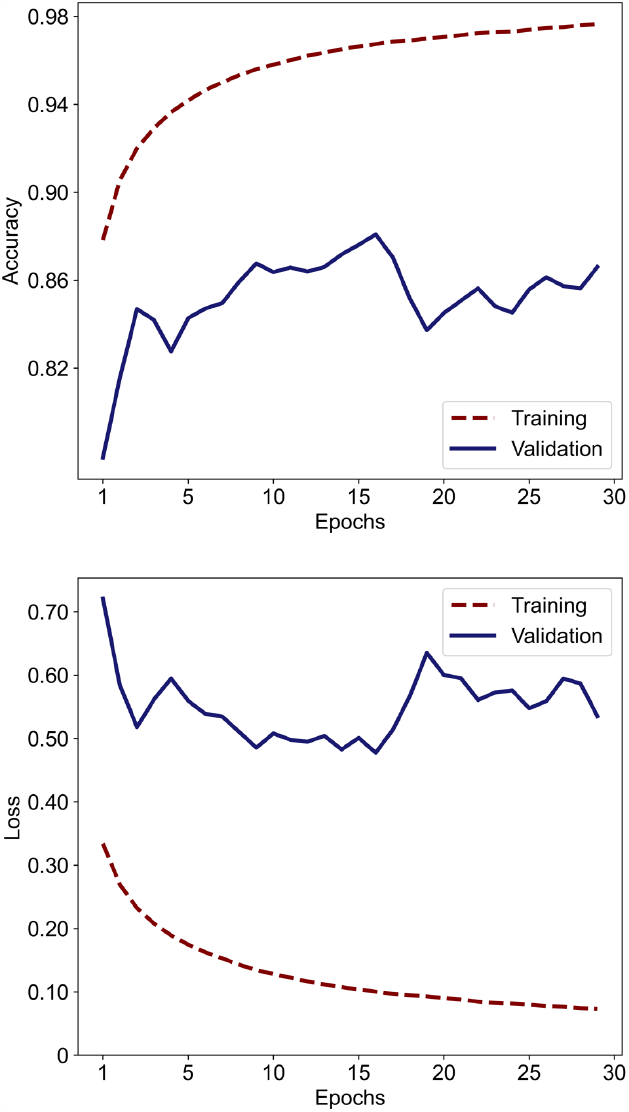
The loss and classification accuracy on the training (maroon) and validation (blue) sets.

## IV. Discussion

In this study, we developed a novel pair of AI-powered smart glasses that uniquely integrates the sensing and deep learning computation for visual perception of human-robot walking environments with high accuracy and low latency. We designed the system using a Raspberry Pi Pico microcontroller and an ArduCam HM0360 camera, which interfaces with the eyeglass frames using 3D-printed mounts that we custom-designed. We then trained and optimized a lightweight and effi-cient convolutional neural network using a MobileNet backbone [22] to classify the walking terrain as either indoor surfaces, outdoor surfaces (grass and dirt), or outdoor surfaces (paved) using over 62,500 images that we manually re-labelled from the Meta Ego4D open-source dataset [21]. Our system could accurately predict complex walking terrains in 1.5 seconds and with high accuracy (∼94%). These results demonstrate, for the first time, the potential to develop fast and accurate egocentric visual perception systems for human-robot walking using embedded computing.

Compared to previous work that required off-device inference due to limited computational power and the high memory demands of machine learning models, our system leverages the latest advances in microcontroller hardware, efficient neural network architectures, and model compression algorithms in order to develop a novel integrated system. As a result, our smart glasses can uniquely process and classify images of walking environments with low latency without a dependency on inconsistent wireless communication to desktop machines or cloud computing.

Compared to leg, waist, and chest-mounted systems [4]-[11], our smart glasses offer several benefits due to its human-centered design. Our system was trained and tested using egocentric images, also known as first-person vision. This point-of-view mimics the biological vision system and takes into consideration the orientation of the user’s head, which has practical implications for inferring intent. Additionally, our smart glasses do not have explicit requirements for the pose or viewing angles compared to other systems [23], which rely on manual heuristics and rule-based thresholds for both the users and environments.

Despite this progress, our study still has several limitations. Our inference speed should be further improved for real-time control. This would be especially important for rehabilitation robotics like exoskeletons that need to dynamically adapt online to different walking terrains. Another area for improvement would be to increase the number of environmental states in our classification model, therein allowing our system to be more applicable to a wider range of applications such as providing sensory feedback to persons with visual impairments and autonomous driving with powered wheelchairs. Future research should also consider further miniaturization of the mechatronic components in order to improve user comfort and acceptance. While our MobileNet model [22] resulted in good performance in terms of classification accuracy and embedded inference speed, we also plan to experiment with and compare other lightweight and efficient architectures (e.g., FastViT [24] by Apple or EfficientViT [25] by Microsoft Research) that are suitable for real-time edge computing.

It is important to note that our visual perception system is designed to supplement, not replace, the existing intent recognition systems of robotic prosthetic legs and exoskeletons that use mechanical, inertial, and/or EMG data to estimate the current state of the human-robot-environment system [26]. We view computer vision as a means to improve the speed and accuracy of locomotion mode (intent) recognition by minimizing the search space of potential solutions based on the perceived walking environment. Accordingly, sensor fusion methods will need to be studied in the future in order to integrate our smart glasses with existing intent recognition systems used for robot control.

In summary, here we designed and prototyped a novel pair of AI-powered smart glasses that uniquely integrate sensing and deep learning computation for onboard visual perception of human-robot walking environments. Applications of our technology range from autonomous control of robotic leg prostheses and exoskeletons to providing sensory feedback to persons with visual impairments. Moving forward, we plan to further improve the onboard inference speed and miniaturization of the mechatronic components.

## Acknowledgment

We want to thank members of the Bionics Lab, part of the Artificial Intelligence and Robotics in Rehabilitation Team at the KITE Research Institute, Toronto Rehabilitation Institute, for their support, especially A. Garrett Kurbis and Hannah Smegal. This research is dedicated to the people of Ukraine in response to the 2022 Russian invasion.

## References

[1] J. Engel et al., “Project Aria: A new tool for egocentric multi-modal AI research,” arXiv, Oct. 1, 2023.

[2] L. Novo-Torres, J.-P. Ramirez-Paredes and D. J. Villarreal, “Obstacle recognition using computer vision and convolutional neural networks for powered prosthetic leg applications”, in 2019 41st Annual International Conference of the IEEE Engineering in Medicine and Biology Society (EMBC), pp. 3360–3363, Jul. 2019.

[3] R. L. da Silva, N. Starliper, B. Zhong, H. H. Huang, and E. Lobaton, “Evaluation of embedded platforms for lower limb prosthesis with visual sensing capabilities.” arXiv, Jun. 26, 2020.

[4] H. A. Al-Dabbagh and R. Ronsse, “Depth vision-based terrain detection algorithm during human locomotion,” IEEE Trans. Med. Robot. Bionics, vol. 4, no. 4, pp. 1010–1021, Nov. 2022.

[5] K. Karacan, J. T. Meyer, H. I. Bozma, R. Gassert, and E. Samur, “An environment recognition and parameterization system for shared-control of a powered lower-limb exoskeleton,” in 2020 8th IEEE RAS/EMBS International Conference for Biomedical Robotics and Biomechatronics (BioRob), pp. 623–628, Nov. 2020.

[6] G. Khademi and D. Simon, “Convolutional neural networks for environmentally aware locomotion mode recognition of lower-limb amputees,” in 2019 ASME Dynamic Systems and Control Conference, Nov. 2019.

[7] A. G. Kurbis, B. Laschowski, and A. Mihailidis, “Stair recognition for robotic exoskeleton control using computer vision and deep learning,”in 2022 International Conference on Rehabilitation Robotics (ICORR), pp. 1–6, Jul. 2022.

[8] A. G. Kurbis, A. Mihailidis, and B. Laschowski, “Development and mobile deployment of a stair recognition system for human-robot locomotion.” bioRxiv, Apr. 28, 2023.

[9] D. Kuzmenko, O. Tsepa, A. G. Kurbis, A. Mihailidis, and B. Laschowski, “Efficient visual perception of human-robot walking environments using semi-supervised learning.” bioRxiv, Jun. 29, 2023.

[10] Ivanyuk-Skulskiy, A. G. Kurbis, A. Mihailidis, and B. Laschowski, “Sequential image classification of human-robot walking environments using temporal neural networks,” bioRxiv, 2023.

[11] A. G. Kurbis, D. Kuzmenko, B. Ivanyuk-Skulskiy, A. Mihailidis, and A. Laschowski, “StairNet: Visual recognition of stairs for human-robot locomotion,” bioRxiv, 2023.

[12] N. E. Krausz and L. J. Hargrove, “Recognition of ascending stairs from 2D images for control of powered lower limb prostheses,” in 2015 7th International IEEE/EMBS Conference on Neural Engineering (NER), pp. 615–618, Apr. 2015.

[13] V. Rai, D. Boe, and E. Rombokas, “Vision for prosthesis control using unsupervised labeling of training data,” in 2020 IEEE-RAS 20th International Conference on Humanoid Robots (Humanoids), pp. 326–333, Jul. 2021.

[14] Pan et al., “COPILOT: Human-environment collision prediction and localization from egocentric videos,” arXiv, Mar. 26, 2023.

[15] A. Sharma and E. Rombokas, “Improving IMU-based prediction of lower limb kinematics in natural environments using egocentric optical flow,” IEEE Trans. Neural Syst. Rehabil. Eng., vol. 30, pp. 699–708, 2022.

[16] Tricomi et al., “Environment-based assistance modulation for a hip exosuit via computer vision,” IEEE Robot. Autom. Lett., vol. 8, no. 5, pp. 2550–2557, May 2023.

[17] “Google Glass Teardown.” http://www.catwig.com/google-glass-teardown/.

[18] “Discover Ray-Ban Stories Features.” https://www.ray-ban.com/canada/en/discover-rayban-stories/clp.

[19] O. Tsepa, R. Burakov, B. Laschowski, and A. Mihailidis, “Continuous prediction of leg kinematics during walking using inertial sensors, smart glasses, and embedded computing.” in IEEE International Conference on Robotics and Automation (ICRA), Jul. 2023.

[20] “Arducam HM0360 VGA SPI Camera Module for Raspberry Pi Pico,” https://www.arducam.com/product/arducam-hm0360-vga-spi-camera-module-for-raspberry-pi-pico-2/.

[21] K. Grauman et al., “Ego4D: Around the world in 3,000 hours of egocentric video,” in 2022 IEEE/CVF Conference on Computer Vision and Pattern Recognition (CVPR), pp. 18973–18990, Jun. 2022.

[22] A. G. Howard et al., “MobileNets: Efficient convolutional neural networks for mobile vision applications,” arXiv, Apr. 17, 2017.

[23] N. E. Krausz, T. Lenzi, and L. J. Hargrove, “Depth sensing for improved control of lower limb prostheses,” IEEE Trans. Biomed. Eng., vol. 62, no.11, pp. 2576–2587, Nov. 2015.

[24] P. K. A. Vasu et al., “FastViT: A fast hybrid vision transformer using structural reparameterization,” arXiv, Aug. 17, 2023.

[25] X. Liu et al., “EfficientViT: Memory efficient vision transformer with cascaded group attention,” arXiv, May. 11, 2023.

[26] R. Gehlhar, M. Tucker, A. J. Young, and A. D. Ames, “A review of current state-of-the-art control methods for lower-limb powered prostheses,” Annual Reviews in Control, vol. 55, pp. 142–164, 2023.

